# Integrated Analysis Reveals Immunogenic Cell Death in Sepsis-induced Cardiomyopathy

**DOI:** 10.1101/2024.03.08.583644

**Authors:** qinxue wang, haobin huang

## Abstract

**Background:** Sepsis-induced cardiomyopathy (SIC) poses a significant challenge in critical care, necessitating comprehensive understanding and innovative diagnostic approaches. This study explores the immune-related molecular intricacies underlying SIC, employing bioinformatics analyses and machine learning techniques.

**Methods:** RNA-seq and scRNA-seq datasets (GSE79962 and GSE190856) were obtained from the Gene Expression Omnibus (GEO). After initial quality control and preprocessing, scRNA-seq data (GSE190856) were analyzed using the Seurat package, including cell clustering and annotation. The CellChat package was then used to analyze immune cell interactions. Unsupervised clustering of SIC patients was performed based on differentially expressed ICD-related genes (GSE79962). Immune cell infiltration and gene set variation analysis were conducted, and weighted gene co-expression network analysis identified co-expression modules. A predictive signature for SIC was constructed through machine learning methods.

**Results:** Trough analyzing the GSE190856 scRNA-seq dataset, the communication between macrophages/monocytes and lymphocytes was found to be enhanced in mouse myocardial tissue during the early onset of SIC. Meanwhile, the expression level of ICD-related genes was upregulated in the monocytes infiltrating to the heart. These results suggestted that ICD may play a crucial role in the pathogenesis of SIC, which had been verified by the upregulated expression of ICD-related genes in the hearts of SIC patients in the GSE79962 dataset. The SIC patients were classified to 2 clusters, with cluster 1 exhibited an upregulation of the renin-angiotensin system, while cluster 2 displayed heightened activity in the RIG-I-like receptor signaling pathway. After comparing four machine learning models, the support vector machine (SVM) model exhibited better discrimination for SIC patients. By correlating the expression levels of the five crucial genes contained in this model with the clinical features of SIC patients, we found that JARID2 was negatively related to the Left Ventricular Ejection Fractions, while TNIP2 was negatively related to the variety of inotropes and vasopressors used in the SIC patients.

**Conclusion:** This research unveils the correlation between ICD and SIC, offering insights into immune activity in the hearts during sepsis. The constructed SVM model with selected genes provides a promising molecular strategy for SIC diagnosis.

## Introduction

Sepsis, a life-threatening condition triggered by the body’s extreme response to infection, remains a global health challenge with alarmingly high mortality rates. Among the myriad of complications associated with sepsis, sepsis-induced cardiomyopathy (SIC) has emerged as a significant concern[1]. Characterized by reversible myocardial dysfunction and structural alterations, SIC has been the subject of intensive research and clinical investigation.

The vulnerability of the heart during sepsis significantly impacts patient prognosis, with mortality rates exceeding two to three times when cardiac dysfunction is present. Unfortunately, specific diagnostic criteria for SIC are lacking, and currently clinical diagnosis relies on comprehensive assessments of cardiac systolic dysfunction, perfusion abnormalities, and increased biomarkers. Despite its clinical significance, there are currently no effective therapeutic agents or procedures specifically tailored to address SIC. Therefore, a thorough understanding of the mechanisms underlying SIC is highly necessary.

In recent years, molecular biology and bioinformatics technologies have advanced our understanding of the pathophysiology of sepsis and its cardiac complications. Within this context, regulated cell death emerges as one of the most noteworthy research directions. Dapagliflozin, Interferon Regulatory Factor-2 (IRF2), Omega-3 polyunsaturated fatty acids, and Jujuboside A have been identified to mitigate the severity of pathological reactions during SIC by suppressing apoptosis[2–5]. On the other hand, ferritinophagy-mediated ferroptosis has been found to play a crucial role in the pathogenesis of SIC[6]. However, further exploration is required to determine whether additional forms of regulated cell death exist in the pathogenesis of SIC.

In this study, based on the analysis of RNA sequencing data, we will analyze the potential forms of regulated cell death in SIC. Subsequently, through GSVA analysis, WGCNA analysis, and machine learning, we will identify hub genes related to SIC, which may provide new targets for the diagnosis and treatment of SIC.

## Methods

### Data Retrieval for Bioinformatics Analysis

RNA-seq and scRNA-seq datasets related to SIC were obtained from the Gene Expression Omnibus (GEO) database. Following a comprehensive search, two datasets, namely GSE79962 and GSE190856, were chosen for further investigation. The GSE79962 dataset provided bulk RNA sequencing data from human hearts[7], and the GSE190856 dataset offered single-cell RNA sequencing data for immune cells in mouse hearts[8]. For subsequent analyses, Transcripts per Kilobase Million (TPM) values were extracted, and genes with an average expression level below 0.1 were excluded.

### scRNA-seq data process

The scRNA-seq dataset GSE190856 was analyzed using the R package Seurat[9]. To ensure data quality, an initial quality control (QC) step was executed by retaining cells with less than 10% mitochondrial gene content and genes expressed in at least three cells within the expression range of 200 to 7000. We identified a set of highly variable genes for further exploration, with the number set at 2000. To mitigate batch effects among data from different samples, the "Harmony" package was employed. Cell clusters were then generated using the "FindClusters" and "FindNeighbors" functions, and the "t-SNE" method was utilized for cluster visualization. Cell annotation was performed based on marker genes associated with different cell types. Cell to cell contact among different immune cells was analyzed and visualized using the R package CellChat[10]. The expression levels for ICD-related genes among the immune cells were calculated using the "AddModuleScore" function within the Seurat package.

### Unsupervised clustering of SIC patients

Unsupervised clustering of SIC patients was initiated with the acquisition of differentially expressed ICD-related genes in SIC patients compared with control patients in the GSE79962 dataset. By utilizing the "ConsensusClusterPlus" R package, the SIC patients were allocated to distinct clusters using the k-means algorithm with 1,000 iterations[11]. To determine the optimal cluster number, a maximum subtype value of k (k = 9) was chosen, and a comprehensive evaluation was conducted based on the cumulative distribution function (CDF) curve, consensus matrix, and consistent cluster score.

### Assessing the immune cell infiltration in different ICD related clusters

The GSE190856 dataset has clearly demonstrated the infiltration of various immune cells in heart during SIC. To explore the relationship between ICD related clusters and immune cell infiltration in cardiac tissue during SIC, we utilized the CIBERSORT algorithm to assess the relative abundances of 22 immune cell types in the GSE79962 dataset[12].

### Gene set variation analysis (GSVA) analysis

By using the R package "GSVA", enrichment analysis was performed to explore distinctions in enriched gene sets between different clusters[13]. The "c2.cp.kegg.symbols" and "c5.go.bp.symbols" were retrieved from the MSigDB database for subsequent GSVA analysis[14]. The "limma" package was employed to discern differentially expressed pathways and biological functions between ICD related clusters by comparing GSVA scores[15].

### Weighted gene co-expression network analysis (WGCNA)

The WGCNA analysis was performed to identify co-expression modules by using the R package "WGCNA"[16]. An optimal soft threshold was chosen for constructing a scale-free network. Subsequently, we converted the weighted adjacency matrix into a topological overlap matrix (TOM) and computed the dissimilarity (dissTOM). Gene clustering and module identification were conducted using the Dynamic Tree Cut approach.

### Construction of predictive signature by integrating machine learning methods

We intersected the genes within the module associated with SIC and the module associated with ICD clustering, as identified by the WGCNA analysis. The intersected genes were considered to be both associated with ICD and SIC. To identity predictive signature with high accuracy, we established machine learning model by using the "caret" R package. The samples in GSE79962 dataset were randomly divided into a training set (70%) or a validation set (30%). Four machine learning models were incorporated, including random forest model (RF), support vector machine model (SVM), generalized linear model (GLM), and eXtreme Gradient Boosting (XGB)[17, 18]. The "DALEX" package was used to interpret and visualize the residual distribution and feature importance among the four machine learning models. On this basis, we determined the optimal machine learning model and the top five crucial variables were regarded as the key predictive genes linked to SIC.

## Results

### Enhanced Communication Between Macrophages and Lymphocytes in SIC

The GSE190856 dataset comprises scRNA-seq data of immune cells in the mouse hearts at steady state and on days 3/7/21 after cecal ligation and puncture (CLP). Immune cells in the dataset were annotated based on the expression of characteristic molecules. A total of six cell clusters were identified in the SIC hearts, namely macrophages (Adgre1, Fcgrt, Timd4, Retnla, Lyve1, Cd163, Folr2, CCR2, H2-Aa, and H2-Eb1), monocytes (Plac8, Fn1, Ace, Itgal, Napsa, Gngt2, and Chil3), neutrophils (S100a8/a9, Retnlg, Ifitm1, Lcn2, Ngp, and Hp), natural killer (NK)/T cells (Cd3e, Klrk1, Ccl5, Gzma, Il1rl1, Gata3, and Il7r), B cells (Igkc, Ly6d, Ebf1, Cd79b, Ms4a1, and Cd79a) and cycling cells (Mki67, Stmn1, Mki67, Top2a, Ube2c, and Stmn1). Macrophages constituted the predominant immune cell type. However, a decline in macrophage percent was noted on day 3 post-CLP, followed by an increase on days 7 and 21. While monocyte infiltration was negligible under steady-state conditions, our findings revealed an influx of monocytes on day 3 after CLP, reaching a peak on day 7 and gradually decreasing by day 21 (Figure 1A). Due T/B cell markers were also expressed in the cycling cells cluster, we considered the cluster as lymphocytes in a rapid proliferative state. CellChat analysis suggested that cell to cell contact between macrophages/monocytes and lymphocytes were enhanced after CLP, indicating increased cell interactions between them in the early pathogenesis of SIC (Figure 1B).

### The expression profile of ICD-related genes in different immune cells during SIC

The enhanced communication between macrophages/monocytes and lymphocytes has led us to hypothesize the involvement of a form of regulated cell death, immunogenic cell death (ICD), in the pathogenesis of SIC. To assess the extent of immunogenic cell death (ICD) activity across distinct immune cell types, we further analyzed the GSE190856 scRNA-seq dataset by using the "AddModuleScore" function within the Seurat package, enabling the computation of expression levels for ICD-related genes across all cells. We observed high ICD activity in the monocytes infiltrating to the heart after CLP (Figure 2).

### Expression of ICD-related genes in the hearts of SIC patients

The above results suggested that ICD plays a significant role in animal models of SIC. We further aim to validate whether it plays an equally important role in SIC patients. Through literature review, we identified 34 genes associated with ICD (Supplementary Table 1). Utilizing the GSE79962 dataset, we assessed the expression of these genes in the hearts of SIC patients. The results revealed a significant upregulation in the expression of IFNGR1, IL1R1, IL6, LY96, EIF2AK3, MYD88, and IL17RA and a marked downregulation in the expression of HSP90AA1, IFNG, and PRF1 when compared with the control group (Figure 3A). Next, we examined the regulatory relationships among these differentially expressed ICD-related genes. The results revealed strong regulatory connections between IFNGR1 and IL6, HSP90AA1 and PRF1, LY96 and IL1R1, as well as IL1R1 and MYD88 (Figure 3B).

### Identifying ICD-Associated Clusters in SIC

To unravel expression patterns of the 10 differentially expressed ICD-related genes (ICDRGs) in SIC, we employed a consensus clustering algorithm to categorize the 20 SIC patients based on the expression profiles of ICDRGs. The optimal stability of cluster numbers was achieved when setting the k value to two (k = 2, Figure 4A). Notably, the consistency score reached its peak when k = 2 (Figure 4B). Compared to the C2 cluster, the expression level of IL6 and IFNGR1 was higher in the C1 cluster (Figure 4C). Principal component analysis (PCA) underscored a significant distinction between these two clusters (Figure 4D).

### Immune infiltration characteristics between ICD clusters

In the previous section, we observed monocytes infiltration in the hearts of the SIC mice model, and these cells exhibited strong ICD activity. We further explored the characteristics of immune cell infiltration in the hearts of SIC patients from the two ICD sub-clusters by using the CIBERSORT algorithm. The results revealed a more pronounced infiltration of activated mast cells in patients from cluster 1 (Figures 5A、B).

### Gene Set Variation Analysis between ICD clusters

We employed GSVA analysis to delve deeper into the functional disparities between the two clusters. The C2 cluster exhibited enhanced regulation of cellular processes, including chromatin organization, protein modification, and biosynthetic pathways, which suggested a dynamic and multifaceted cellular response during SIC. On the other hand, the C2 cluster exhibited suppressed activities related to cardiac muscle cell action potential, lipid metabolism, extracellular matrix regulation, dendritic cell differentiation, and various metabolic processes when comparing with the C1 cluster (Figure 6A). Next, we utilized the KEGG database to analyze the enriched pathways in these two clusters. The results revealed that the C1 cluster exhibited higher activity in regulating cardiac rhythm and vascular contraction, cell adhesion, antigen processing and presentation, and the regulation of various signaling pathways (Figure 6B). These findings suggested that the C1 cluster may undergo more adaptive immune processes.

### Identifying the hub gene modules associated with SIC and ICD clusters

To unveil the hub gene module in SIC, we employed a WGCNA pipeline to construct co-expression modules for both control and SIC subjects. The top 25% genes with the highest variance were selected for further analysis. To build a scale-free network, the soft-threshold was set at seven based on the assessment of scale-free index and mean connectivity at different powers (Figure 7A). Seven modules were detected using the Dynamic Tree Cut method (Figure 7B). These seven gene modules were then scrutinized for their co-expression with clinical features (Control or SIC). Notably, the turquoise module demonstrated the most robust association with SIC and was selected as the SIC related module (Figure 7C). Genes in this module with gene significance for SIC greater than 0.5 and module membership greater than 0.8 were selected as hub genes for SIC(Figure 7D).

We then continued to use WGCNA to further identify the hub gene module related to ICD clusters. The soft-threshold was set at nine to obtain a stable scale-free index and mean connectivity when building a scale-free network (Figure 8A). Eleven modules, detected using the Dynamic Tree Cut method (Figure 8B), were then inspected for their co-expression with ICD clusters (C1 or C2). Remarkably, the purple module exhibited the strongest correlation with ICD clusters (Figure 8C). Thirteen genes in this module, including SH3BGRL, RELB, NFKB2, CAMK1D, SCN5A, LAMA4, CCL8, NFKBIA, EMCN, CSF3, PLXDC2, ICAM1, and CXCL1, with gene significance greater than 0.5 and module membership greater than 0.8, were considered to be significantly associated with ICD clustering (Figure 8D).

### Identifying the gene sets for the diagnosis of SIC using machine learning models

Finally, to identify gene sets with high diagnostic value for SIC, we employed four machine learning models, that were random forest model (RF), support vector machine model (SVM), generalized linear model (GLM), and eXtreme Gradient Boosting model(XGB). After the former WGCNA analysis, we identified gene modules related to SIC and ICD. By intersecting the two modules, 88 genes were obtained for evaluation in the machine learning models (Figure 9A). Through training and validation of machine learning models, we observed that the SVM model exhibited the lowest residual among the four models (Figures 9B/C). Furthermore, by plotting ROC curves in the validation set for each model, we observed that the area under the ROC curve (AUC) for the SVM model approached 1. Therefore, the SVM model may offer better discrimination for SIC patients. Subsequently, we ranked the variables within the SVM model based on root mean square error (RMSE) values and selected the top 5 variables (SULF2, TNIP2, GABARAPL1, JARID2, TMEM173) to form a predictive gene set for SIC (Figure 9D). Finally, we conducted a correlation analysis between the expression levels of these five genes in the hearts of SIC patients and their clinical characteristics. The results revealed a negative correlation between the expression levels of JARID2 and the Left Ventricular Ejection Fractions (LVEF), meanwhile, elevated TNIP2 expression was associated with a decreased variety of inotropes and vasopressors used in the SIC patients (Figure 9E). Additionally, we also found that the expression level of JARID2 and TNIP2 in cluster 1 was significantly higher than that in cluster 2.

## Discussion

In this study, we first explored immune cell interactions during SIC by secondary analysis of a mouse single-cell RNA sequencing dataset. We found enhanced communication between macrophages/monocytes and lymphocytes in the heart tissue during SIC and up-regulation of ICDRGs in myocardial-infiltrating monocytes, suggesting that immune activation and ICD may play an important role in the development of SIC. We further validated the up-regulation of ICDRGs on myocardial transcriptomic data of SIC patients and then classified them into two categories based on ICDRGs features to elucidate their heterogeneity. SIC and ICD related gene modules were identified by incorporating the WGCNA pipeline. Finally, machine learning models were employed to identify a predictive gene set for SIC. After comparing four machine learning models, we selected the SVM model to construct a molecular diagnostic model for SIC based on five genes (SULF2, TNIP2, GABARAPL1, JARID2, and TMEM173), which provides a new molecular strategy for the diagnosis of SIC.

The pathogenesis of SIC involves multiple aspects such as inflammation, oxidative stress, lipid peroxidation, mitochondrial dysfunction, and energy metabolism disorders[1]. Due to its complex and not fully elucidated mechanisms, clinical diagnosis lacks specific biomarkers and consistent criteria, meanwhile, there are currently no effective therapeutic drugs specifically targeting SIC[1, 19]. In recent years, with the advancement of molecular biology, bioinformatics analysis, and other technologies, our understanding of the pathological processes of SIC has deepened. Beyond clinical phenotypes, we can categorize the SIC patients on the basis of different molecular characteristics, and targeted molecular therapies can be developed accordingly. This may be beneficial in enhancing the therapeutic efficacy and improving the prognosis of SIC patients.

In this study, based on the enhanced communication between macrophages/monocytes and lymphocytes as well as differentially expressed ICDRGs in the SIC patients’ hearts, we inferred that ICD has played a crucial role in the pathogenesis of SIC. ICD is a specific variant of regulatory cell death driven by stress, which can cause immunogenic cell death under conditions with concurrent antigenicity, adjuvanticity, and a suitable microenvironment[20, 21]. ICD has been shown to play a positive role in tumor immunity, and various treatments for tumors have been shown to positively activate ICD, exerting anti-tumor effects through the immune process[22]. When the body is infected by pathogens, cells undergo immunogenic stress and release endogenous damage-associated molecular patterns (DAMPs), which, along with conserved components of bacteria and viruses (e.g., lipopolysaccharides and double-stranded RNA), form microbe-associated molecular patterns (MAMPs). The MAMPs are collectively sensed by pattern recognition receptors (PRRs) expressed by bone marrow cells, thus inducing the occurrence of ICD[23]. However, it is still unclear whether ICD plays a beneficial or harmful role in SIC.

In this study, through the analysis of the single-cell RNA sequencing dataset GSE190856, we clarified the characteristics of immune cell infiltration in myocardial tissues during SIC. Cell to cell contacts between macrophages/monocytes and lymphocytes during the early stage of SIC were demonstrated to be enhanced, moreover, the ICD activity of monocytes infiltrating to the heart was high, indicating the involvement of ICD in the development of SIC. We then comprehensively analyzed the expression profiles of 34 ICD-related genes in the myocardium of SIC patients and control groups, and found that compared to the control group, the expression of multiple ICD-related genes were significantly changed, suggesting the importance of ICD in the pathogenesis of SIC. We further analyzed the regulatory interactions among differentially expressed ICD-related genes, revealing multiple gene groups with regulatory connections, confirming the existence of interactions among ICD-related genes in SIC. Based on these findings, we believe that ICD plays a crucial role in the pathogenesis of SIC and could serve as a target for the diagnosis and treatment of SIC.

This study further divided SIC patients into two clusters based on different ICD characteristics. Cluster 1 had higher levels of IL-6 and IFNGR1 expression, and activated mast cell infiltration was more pronounced. Results from GO and KEGG analyses suggested a more positive cardiovascular system response to septic stimuli for this cluster, manifested by the upregulation of the renin-angiotensin system activity, better maintenance of vascular constriction, heart rate, and myocardial contraction status, and increased perfusion pressure. on the other hand, for the patients in cluster 2, the RIG-I-like receptor (RLR) signaling pathway was more upregulated. This pathway is associated with innate immunity and is involved in the recognition of cytoplasmic dsRNA, which is associated with the subsequent activation of STING, IRF, and NF-κB pathways[24]. During these processes, more protein modifications may occur than that in cluster 1.

In recent years, with the development of artificial intelligence, research on the application of machine learning models in the diagnosis and treatment of sepsis has been increasing. Kim et al. demonstrated that by using an artificial intelligence algorithm (SERA algorithm), the likelihood of early sepsis detection can be increased by 32%, thus facilitating timely treatment and improved prognosis[25]. Liu et al. developed the T4 algorithm, a novel approach for estimating individual treatment effects in time-to-treatment antibiotic stewardship for sepsis, demonstrating superior performance compared to existing models[26]. In our study, we used four machine learning models to predict SIC based on the SIC and ICD related gene modules. We found that the SVM model had the lowest residual error and an AUC closed to 1, indicating that the model had good predictive performance for SIC.

Furthermore, we selected five important variables (SULF2, TNIP2, GABARAPL1, JARID2, TMEM173) from the SVM model to construct a predictive gene set for SIC. SULF2, or sulfatase 2, is a protease that specifically hydrolyzes sulfate ester bonds in the extracellular matrix, mainly acting on 6-O-glucosamine sulfate chains in heparan sulfate glycoproteins (HSPGs)[27]. Currently, most research on SULF2 is related to cancer, which has a promoting effect on various cancers and can also be used as a biomarker for tumor diagnosis and prognosis evaluation[28, 29]. Although there is currently no research on the relationship between SULF2 and sepsis, studies do have shown that SULF2 can promote macrophage M2 polarization, affect TNF-induced inflammatory activation, and upregulate GDF15 expression, all of which may be relevant to the occurrence and development of SIC. Therefore, SULF2 may be a potential intervention target for SIC[30–32]. TNIP2, or TNF-α-induced protein 3-interacting protein 2, is a pivotal protein in the NF-κB network, and multiple components of the NF-κB signaling pathway can interact with it. TNIP2 can inhibit NF-κB activation by blocking the interaction between RIPK1 and its downstream effector IKBKG[33], and it can also exert anti-inflammatory and anti-apoptotic effects through the TLR4/MyD88/NF-κB pathway. In this study, the expression level of TNIP2 was negatively correlated with the categories of vasopressors and inotropes used in SIC patients. Additionally, the expression level of TNIP2 in Cluster 1 was significantly higher than that in Cluster 2, combined with the results of GO and KEGG analysis, it suggests that TNIP2 may play a protective role in SIC[34]. GABARAPL1, or GABA A receptor-associated protein-like 1, is one of the autophagy-related ATG8 family proteins, and is an estrogen-induced gene with relatively high expression in the central nervous system. GABARAPL1 is associated with cell proliferation, invasion, autophagy flux, mitochondrial homeostasis, and cellular metabolism. Decreased expression of GABARAPL1 will lead to reduced autophagy flux and accumulation of damaged mitochondria[35]. Research on the relevance of GABARAPL1 to sepsis and SIC is still lacking, however, as mitochondrial damage is one of the most important mechanisms underlying SIC[1], GABARAPL1 may also play a crucial role in the pathogenesis of SIC and could serve as a therapeutic target for the disease. Jumonji/ARID domain-containing protein 2 (JARID2) is the founding member of the JmjC domain-containing protein family, whose function is mainly related to the regulation of histone methyltransferase activity[36, 37], and is crucial for the cardiac development and epigenetic regulation[38]. Left ventricular overload or failure can induce JARID2 expression[39], and our analysis also suggested that the expression level of JARID2 was negatively correlated with left ventricular contraction function. Therefore, it may serve as a marker for evaluating left ventricular systolic function in SIC. In the present study, the expression level of JARID2 in cluster 2 was significantly lower than that in cluster 1, indicating potentially worse left ventricular function in these patients. These patients closely correspond to SIC patients with relatively worse left ventricular function observed in clinical practice, who may have a poorer prognosis. TMEM173, also known as STING, plays a pivotal role in the cell’s innate immune response to pathogenic cytoplasmic DNA[40]. It is primarily activated by the second messenger cyclic GMP-AMP (cGAMP), and induces type I interferon production through TBK1 and IRF. Studies on STING have shown that it can cause sepsis-related inflammation and organ dysfunction during sepsis. Li et al. demonstrated that STING-IRF3 promotes the development of SIC by activating NLRP3, indicating STING as a potential therapeutic target for SIC[41].

This study has some limitations. First, due to the difficulty in obtaining myocardial tissue samples from SIC patients, the transcriptome data is limited. Currently, there is only one human septic cardiomyopathy transcriptome dataset in the GEO database, and the sample size is limited. Further expansion of the sample size and introducing external validation are needed to further clarify the predictive efficacy of the diagnostic model for SIC. Second, the GSE79962 dataset provided only limited clinical data, restricting the assessment of the correlation between five important genes and clinical indicators. More detailed clinical features are needed in future studies to clarify their correlation with ICD and SIC. Finally, this study is primarily based on comprehensive bioinformatics analysis and lacks additional experimental validation. Therefore, further validation experiments are needed in clinical or animal models.

In conclusion, this study reveals for the first time the correlation between ICD and SIC, classifying patients with different ICD features and elucidating their immune heterogeneity. By using machine learning methods, we also constructed a diagnostic model based on five important genes (SULF2, TNIP2, GABARAPL1, JARID2, TMEM173), providing a new molecular strategy for the diagnosis of SIC. This study opens new directions for understanding the mechanisms and treatment targets of SIC.

---

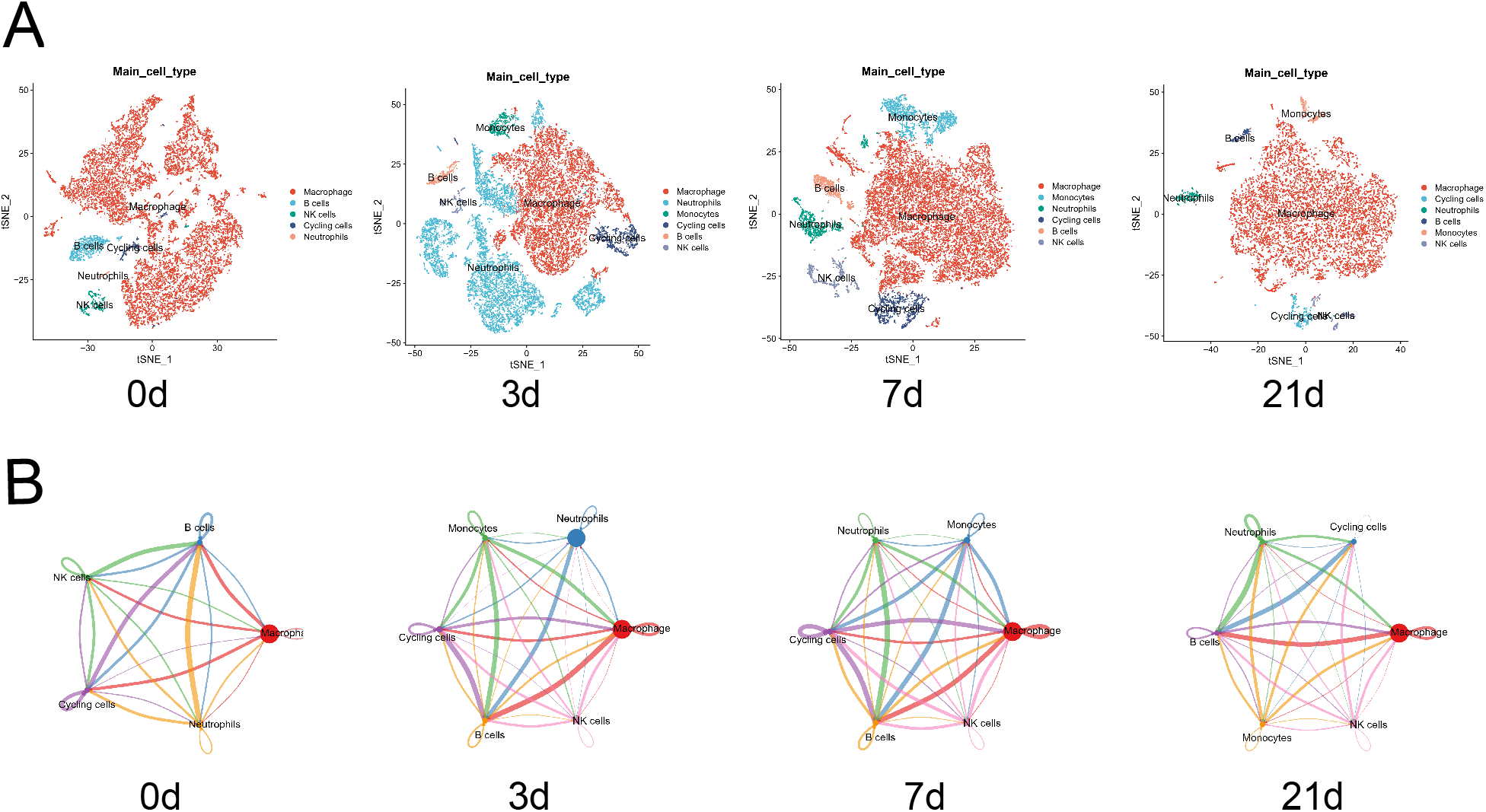

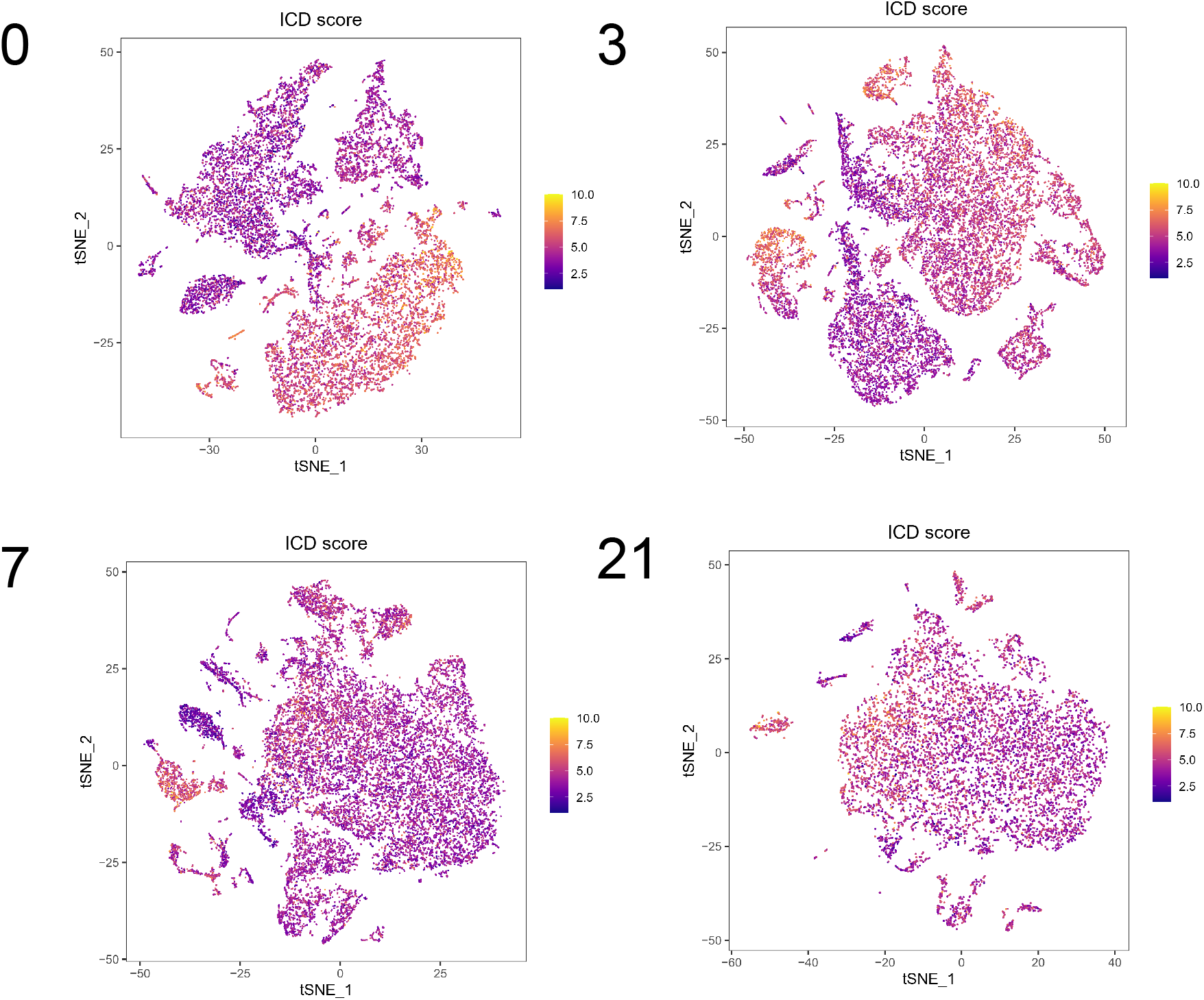

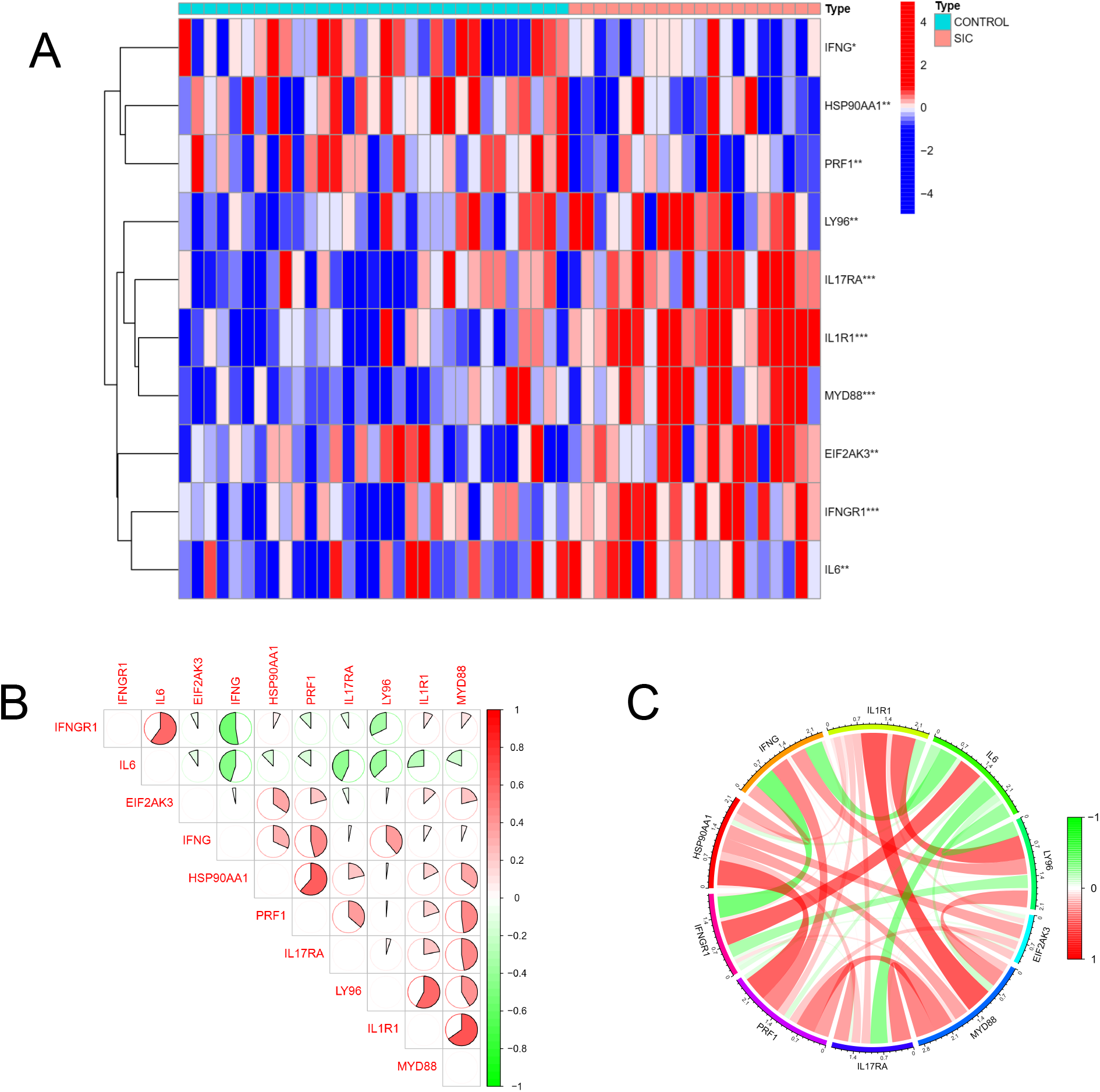

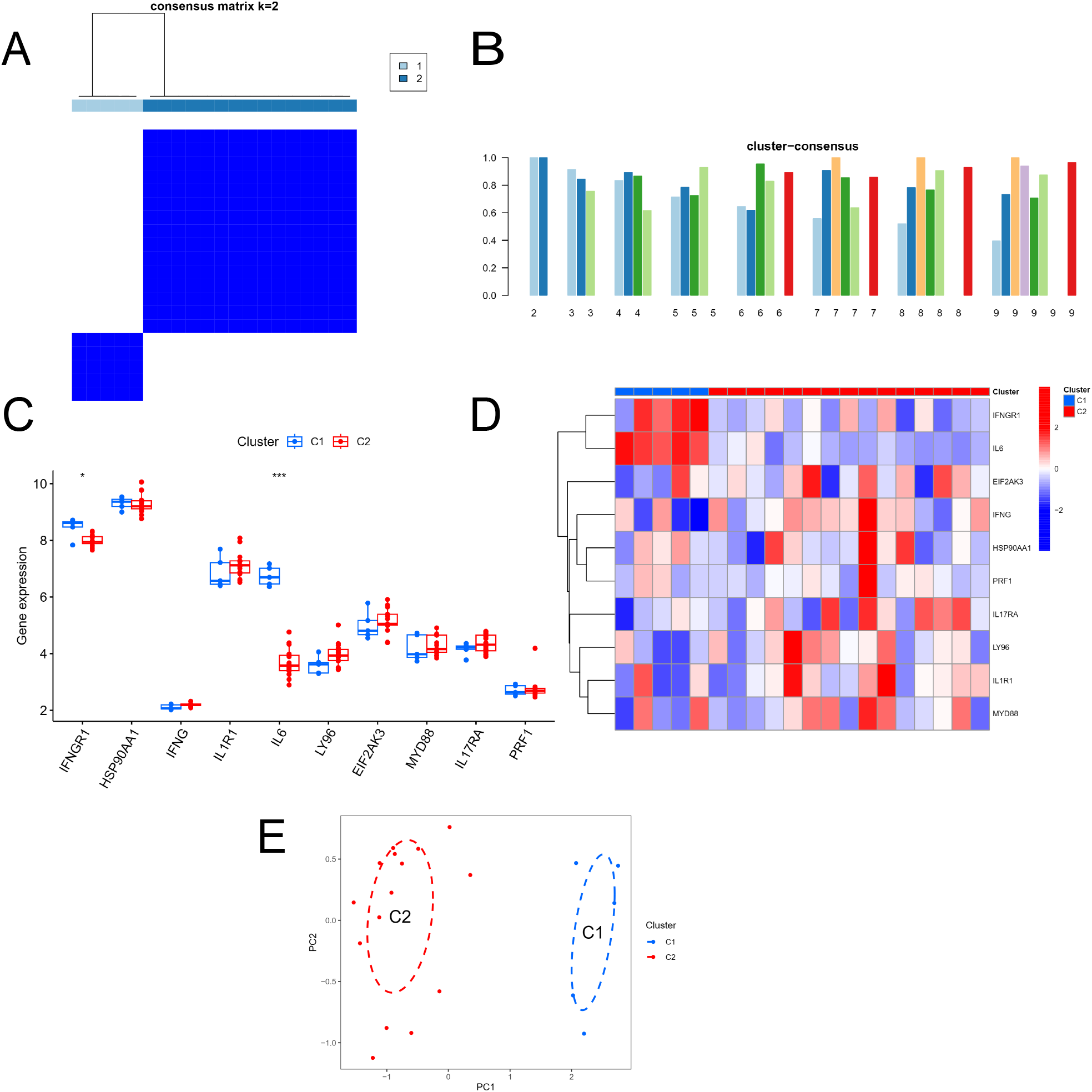

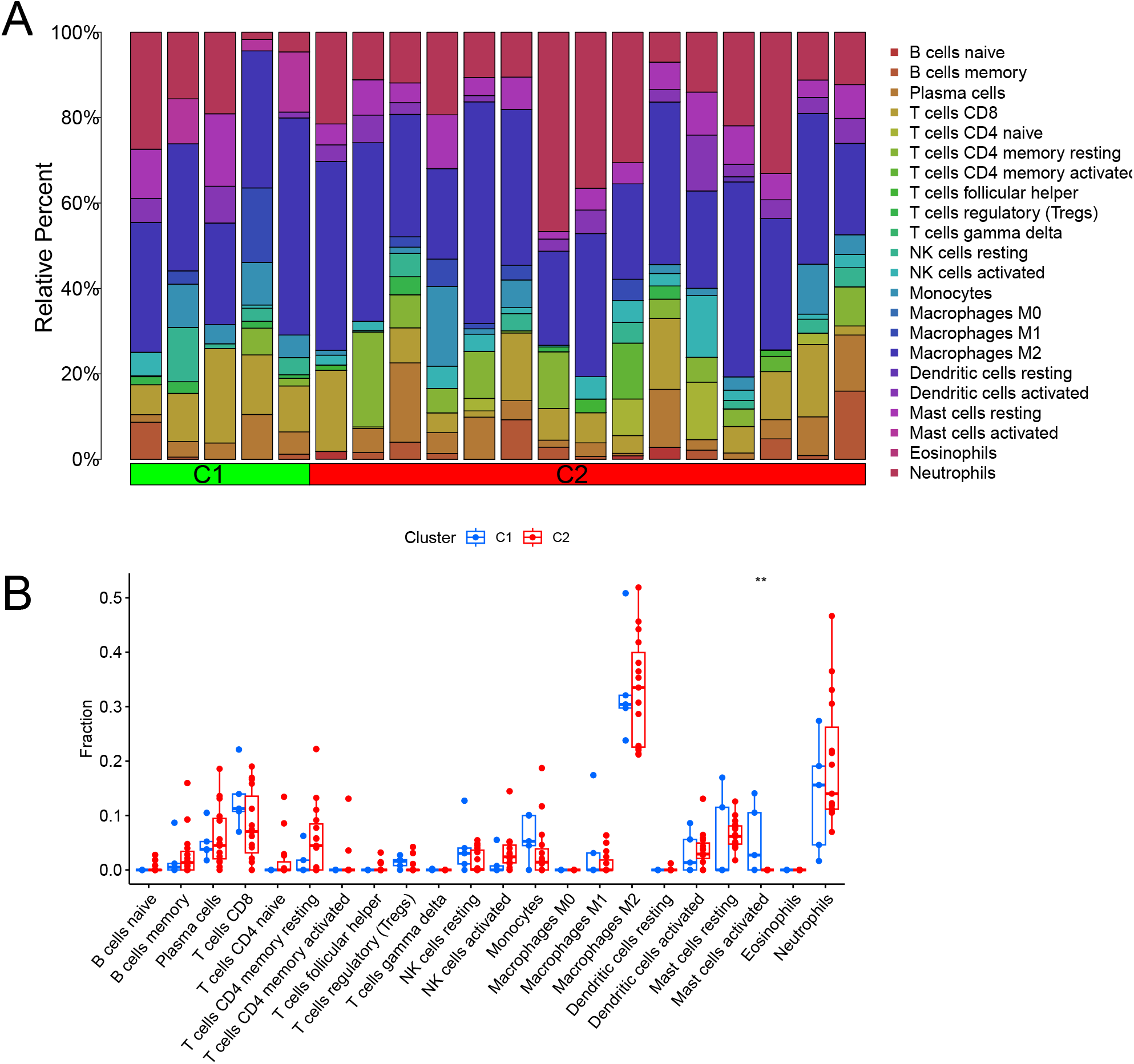

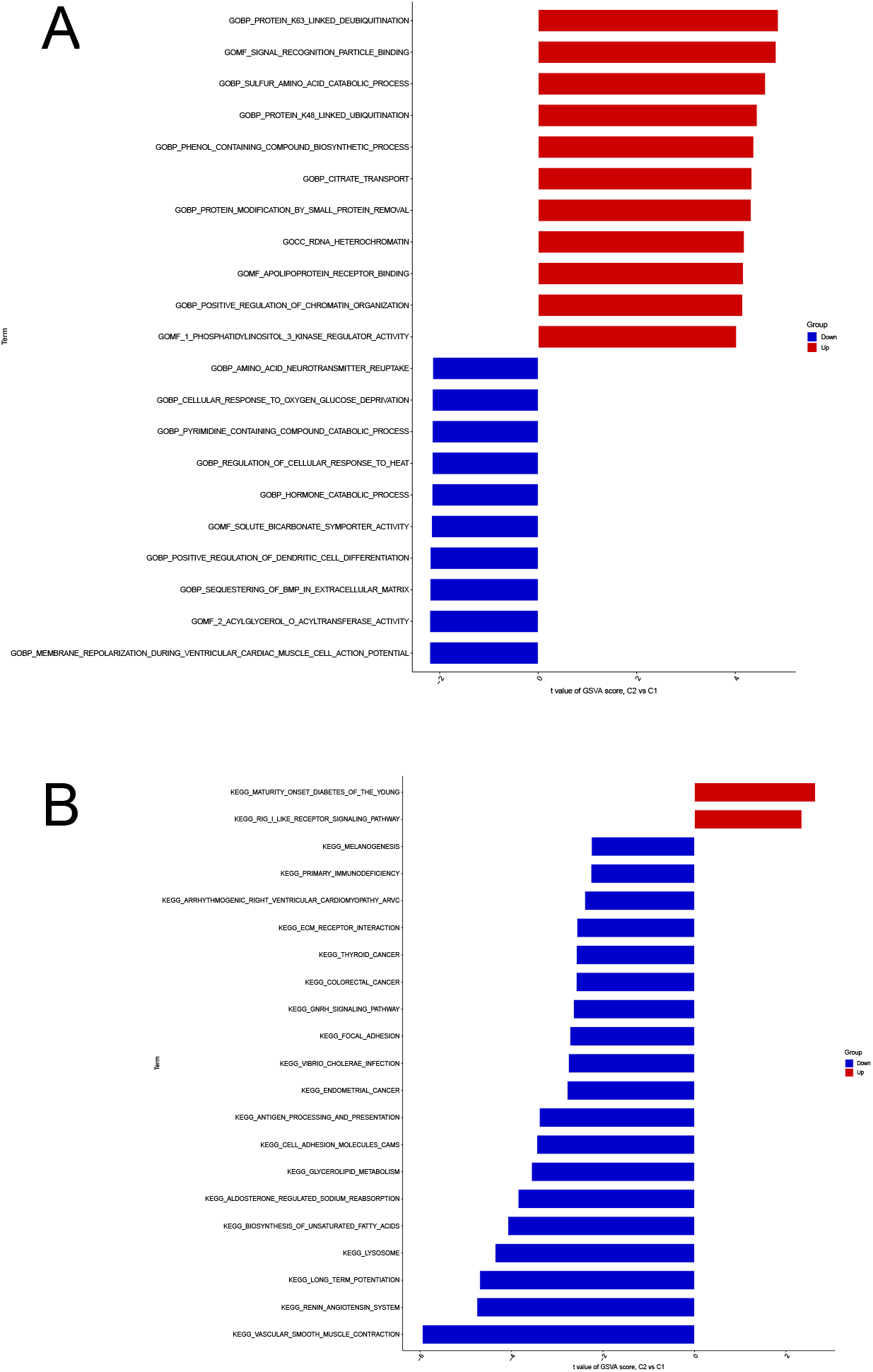

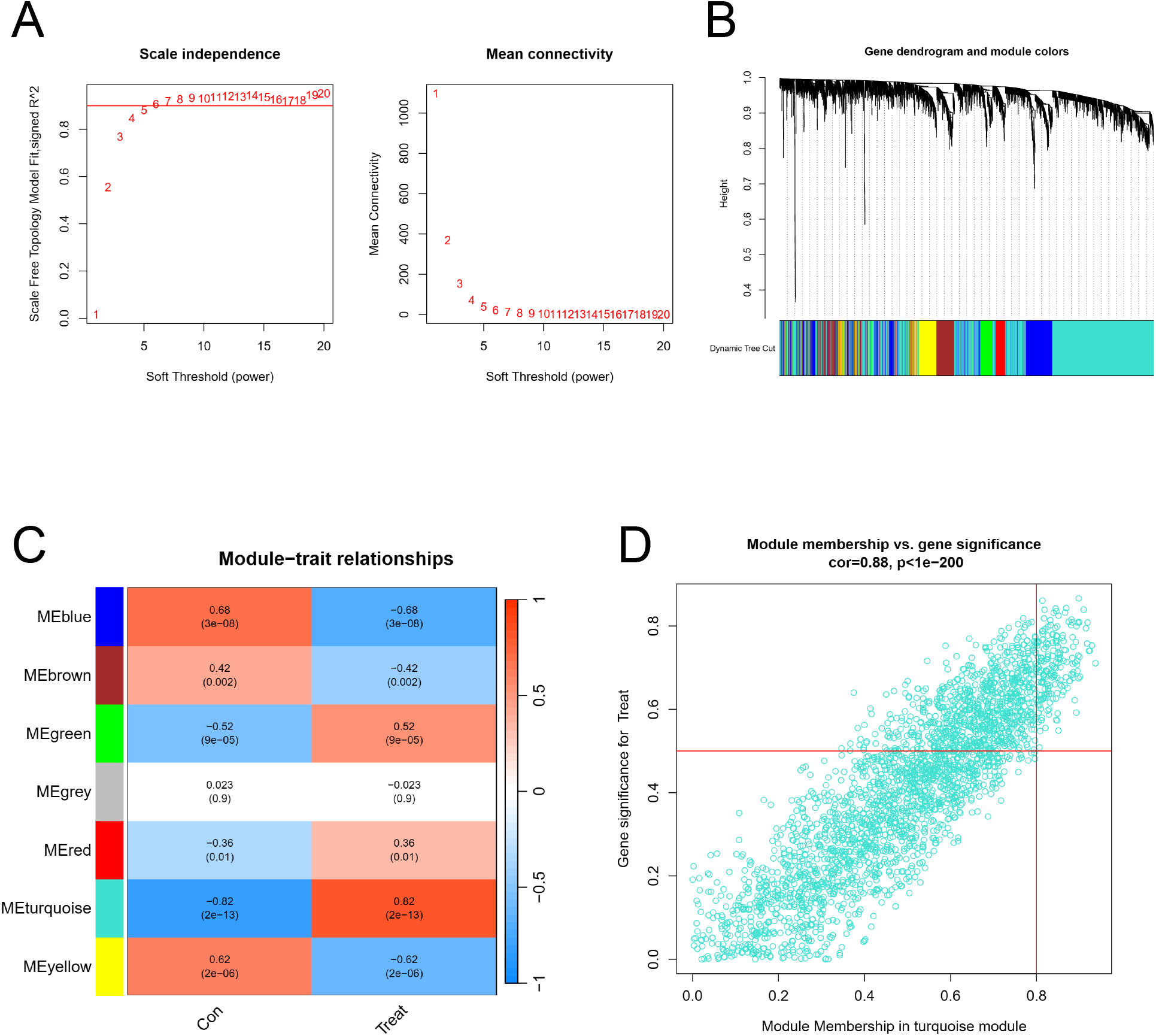

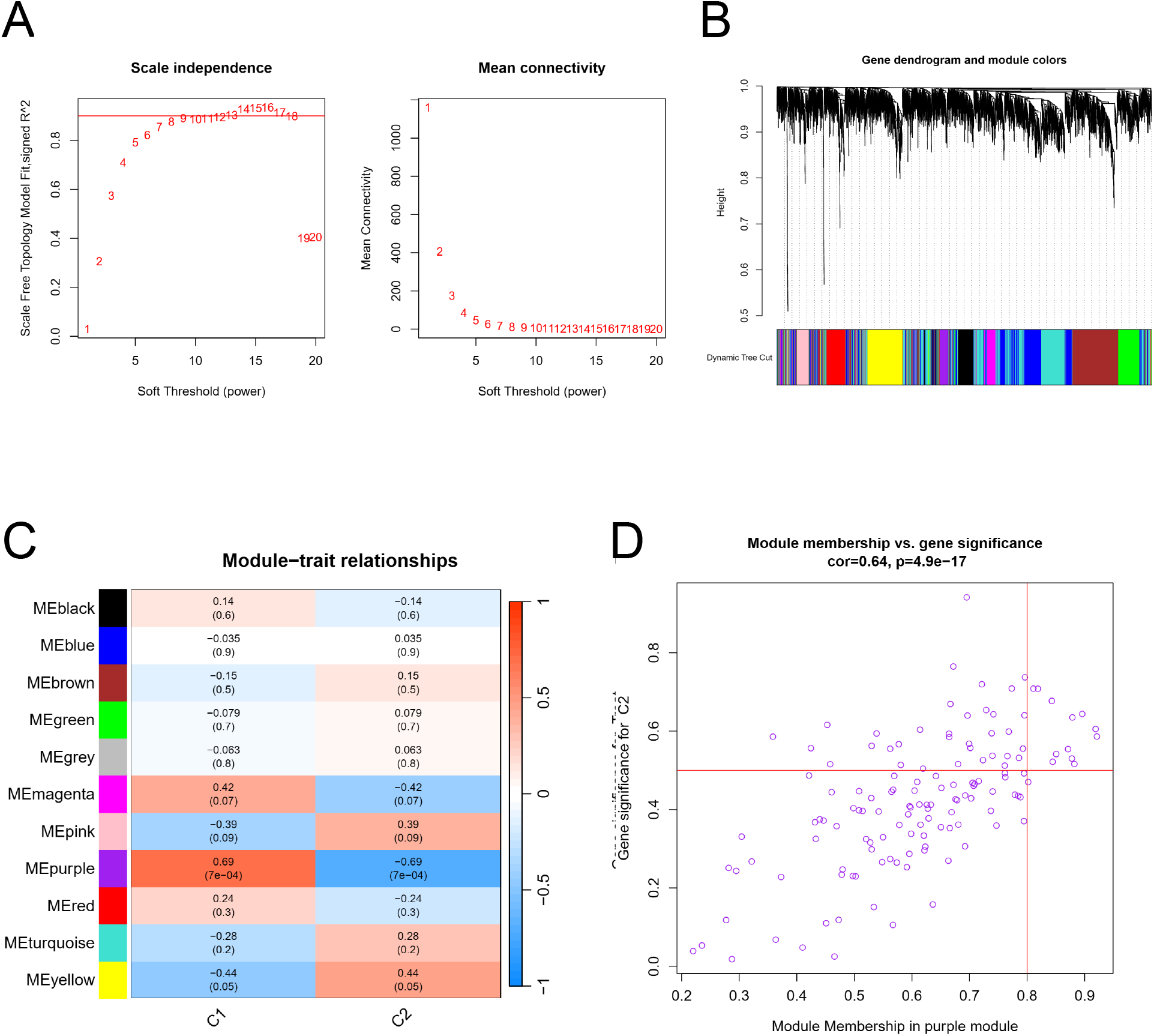

## References

1. Hollenberg SM, Singer M. Pathophysiology of sepsis-induced cardiomyopathy. Nature reviews Cardiology 2021, 18 (6): 424–434.

2. Han X, Liu X, Zhao X, Wang X, Sun Y, Qu C, Liang J, Yang B. Dapagliflozin ameliorates sepsis-induced heart injury by inhibiting cardiomyocyte apoptosis and electrical remodeling through the PI3K/Akt pathway. European journal of pharmacology 2023, 955: 175930.

3. Li T, Luo Q, He L, Li D, Li Q, Wang C, Xie J, Yi C. Interferon Regulatory Factor-2 Binding Protein 2 Ameliorates Sepsis-Induced Cardiomyopathy via AMPK-Mediated Anti-Inflammation and Anti-Apoptosis. Inflammation 2020, 43 (4): 1464–1475.

4. Chen ZS, Yu MM, Wang K, Meng XL, Liu YC, Shou ST, Chai YF. Omega-3 polyunsaturated fatty acids inhibit cardiomyocyte apoptosis and attenuate sepsis-induced cardiomyopathy. Nutrition (Burbank, Los Angeles County, Calif) 2023, 106: 111886.

5. Wang Z, Xiao D, Ji Q, Li Y, Cai Z, Fang L, Huo H, Zhou G, Yan X, Shen L, He B. Jujuboside A attenuates sepsis-induced cardiomyopathy by inhibiting inflammation and regulating autophagy. European journal of pharmacology 2023, 947: 175451.

6. Li N, Wang W, Zhou H, Wu Q, Duan M, Liu C, Wu H, Deng W, Shen D, Tang Q. Ferritinophagy-mediated ferroptosis is involved in sepsis-induced cardiac injury. Free radical biology & medicine 2020, 160: 303–318.

7. Matkovich SJ, Al Khiami B, Efimov IR, Evans S, Vader J, Jain A, Brownstein BH, Hotchkiss RS, Mann DL. Widespread Down-Regulation of Cardiac Mitochondrial and Sarcomeric Genes in Patients With Sepsis. Critical care medicine 2017, 45 (3): 407–414.

8. Zhang K, Wang Y, Chen S, Mao J, Jin Y, Ye H, Zhang Y, Liu X, Gong C, Cheng X, Huang X, Hoeft A, Chen Q, Li X, Fang X. TREM2(hi) resident macrophages protect the septic heart by maintaining cardiomyocyte homeostasis. Nature metabolism 2023, 5(1): 129–146.

9. Butler A, Hoffman P, Smibert P, Papalexi E, Satija R. Integrating single-cell transcriptomic data across different conditions, technologies, and species. Nature biotechnology 2018, 36 (5): 411–420.

10. Jin S, Guerrero-Juarez CF, Zhang L, Chang I, Ramos R, Kuan CH, Myung P, Plikus MV, Nie Q. Inference and analysis of cell-cell communication using CellChat. Nature communications 2021, 12 (1): 1088.

11. Wilkerson MD, Hayes DN. ConsensusClusterPlus: a class discovery tool with confidence assessments and item tracking. Bioinformatics (Oxford, England) 2010, 26 (12): 1572–1573.

12. Newman AM, Liu CL, Green MR, Gentles AJ, Feng W, Xu Y, Hoang CD, Diehn M, Alizadeh AA. Robust enumeration of cell subsets from tissue expression profiles. Nature methods 2015, 12 (5): 453–457.

13. Hänzelmann S, Castelo R, Guinney J. GSVA: gene set variation analysis for microarray and RNA-seq data. BMC bioinformatics 2013, 14: 7.

14. Liberzon A, Birger C, Thorvaldsdóttir H, Ghandi M, Mesirov JP, Tamayo P. The Molecular Signatures Database (MSigDB) hallmark gene set collection. Cell systems 2015, 1(6): 417–425.

15. Ritchie ME, Phipson B, Wu D, Hu Y, Law CW, Shi W, Smyth GK. limma powers differential expression analyses for RNA-sequencing and microarray studies. Nucleic acids research 2015, 43 (7): e47.

16. Langfelder P, Horvath S. WGCNA: an R package for weighted correlation network analysis. BMC bioinformatics 2008, 9: 559.

17. Rigatti SJ. Random Forest. Journal of insurance medicine (New York, NY) 2017, 47 (1): 31–39.

18. Byvatov E, Schneider G. Support vector machine applications in bioinformatics. Applied bioinformatics 2003, 2(2): 67–77.

19. Liu YC, Yu MM, Shou ST, Chai YF. Sepsis-Induced Cardiomyopathy: Mechanisms and Treatments. Frontiers in immunology 2017, 8: 1021.

20. Galluzzi L, Vitale I, Warren S, Adjemian S, Agostinis P, Martinez AB, Chan TA, Coukos G, Demaria S, Deutsch E, Draganov D, Edelson RL, Formenti SC, Fucikova J, Gabriele L, Gaipl US, Gameiro SR, Garg AD, Golden E, Han J, Harrington KJ, Hemminki A, Hodge JW, Hossain DMS, Illidge T, Karin M, Kaufman HL, Kepp O, Kroemer G, Lasarte JJ, Loi S, Lotze MT, Manic G, Merghoub T, Melcher AA, Mossman KL, Prosper F, Rekdal Ø, Rescigno M, Riganti C, Sistigu A, Smyth MJ, Spisek R, Stagg J, Strauss BE, Tang D, Tatsuno K, van Gool SW, Vandenabeele P, Yamazaki T, Zamarin D, Zitvogel L, Cesano A, Marincola FM. Consensus guidelines for the definition, detection and interpretation of immunogenic cell death. Journal for immunotherapy of cancer 2020, 8(1).

21. Kroemer G, Galassi C, Zitvogel L, Galluzzi L. Immunogenic cell stress and death. Nature immunology 2022, 23 (4): 487–500.

22. Fucikova J, Kepp O, Kasikova L, Petroni G, Yamazaki T, Liu P, Zhao L, Spisek R, Kroemer G, Galluzzi L. Detection of immunogenic cell death and its relevance for cancer therapy. Cell death & disease 2020, 11 (11): 1013.

23. Galluzzi L, Buqué A, Kepp O, Zitvogel L, Kroemer G. Immunogenic cell death in cancer and infectious disease. Nature reviews Immunology 2017, 17 (2): 97–111.

24. Arnoult D, Soares F, Tattoli I, Girardin SE. Mitochondria in innate immunity. EMBO reports 2011, 12 (9): 901–910.

25. Goh KH, Wang L, Yeow AYK, Poh H, Li K, Yeow JJL, Tan GYH. Artificial intelligence in sepsis early prediction and diagnosis using unstructured data in healthcare. Nature communications 2021, 12 (1): 711.

26. Liu R, Hunold KM, Caterino JM, Zhang P. Estimating treatment effects for time-to-treatment antibiotic stewardship in sepsis. Nature machine intelligence 2023, 5(4): 421–431.

27. Hanson SR, Best MD, Wong CH. Sulfatases: structure, mechanism, biological activity, inhibition, and synthetic utility. Angewandte Chemie (International ed in English) 2004, 43 (43): 5736–5763.

28. Carr RM, Romecin Duran PA, Tolosa EJ, Ma C, Oseini AM, Moser CD, Banini BA, Huang J, Asumda F, Dhanasekaran R, Graham RP, Toruner MD, Safgren SL, Almada LL, Wang S, Patnaik MM, Roberts LR, Fernandez-Zapico ME. The extracellular sulfatase SULF2 promotes liver tumorigenesis by stimulating assembly of a promoter-looping GLI1-STAT3 transcriptional complex. The Journal of biological chemistry 2020, 295 (9): 2698–2712.

29. Huang J, Li C, Zhang W, Yang F, Wang R, Zhang J, Li W, Yao X. SULF2 is a novel diagnostic and prognostic marker for high-grade bladder cancer with lymphatic metastasis. Annals of translational medicine 2021, 9(18): 1439.

30. Zhang W, Yang F, Zheng Z, Li C, Mao S, Wu Y, Wang R, Zhang J, Zhang Y, Wang H, Li W, Huang J, Yao X. Sulfatase 2 Affects Polarization of M2 Macrophages through the IL-8/JAK2/STAT3 Pathway in Bladder Cancer. Cancers 2022, 15 (1).

31. Onuora S. Sulf2 mediates the effects of TNF in RASFs. Nature reviews Rheumatology 2022, 18 (11): 613.

32. He R, Shi J, Xu D, Yang J, Shen Y, Jiang YS, Tao L, Yang M, Fu X, Yang JY, Liu D, Huo Y, Shen X, Lu P, Niu N, Sun YW, Xue J, Liu W. SULF2 enhances GDF15-SMAD axis to facilitate the initiation and progression of pancreatic cancer. Cancer letters 2022, 538: 215693.

33. Banks CA, Boanca G, Lee ZT, Eubanks CG, Hattem GL, Peak A, Weems LE, Conkright JJ, Florens L, Washburn MP. TNIP2 is a Hub Protein in the NF-κB Network with Both Protein and RNA Mediated Interactions. Molecular & cellular proteomics : MCP 2016, 15 (11): 3435–3449.

34. Yan Z, Chen Y, Zhang X, Hua L, Huang L. Neuroprotective Function of TNFAIP3 Interacting Protein 2 Against Oxygen and Glucose Deprivation/Reoxygenation-Induced Injury in Hippocampal Neuronal HT22 Cells Through Regulation of the TLR4/MyD88/NF-κB Pathway. Neuropsychiatric disease and treatment 2021, 17: 2219–2227.

35. Boyer-Guittaut M, Poillet L, Liang Q, Bôle-Richard E, Ouyang X, Benavides GA, Chakrama FZ, Fraichard A, Darley-Usmar VM, Despouy G, Jouvenot M, Delage-Mourroux R, Zhang J. The role of GABARAPL1/GEC1 in autophagic flux and mitochondrial quality control in MDA-MB-436 breast cancer cells. Autophagy 2014, 10 (6): 986–1003.

36. Klose RJ, Kallin EM, Zhang Y. JmjC-domain-containing proteins and histone demethylation. Nature reviews Genetics 2006, 7(9): 715–727.

37. Cloos PA, Christensen J, Agger K, Helin K. Erasing the methyl mark: histone demethylases at the center of cellular differentiation and disease. Genes & development 2008, 22 (9): 1115–1140.

38. Cho E, Mysliwiec MR, Carlson CD, Ansari A, Schwartz RJ, Lee Y. Cardiac-specific developmental and epigenetic functions of Jarid2 during embryonic development. The Journal of biological chemistry 2018, 293 (30): 11659–11673.

39. Bovill E, Westaby S, Reji S, Sayeed R, Crisp A, Shaw T. Induction by left ventricular overload and left ventricular failure of the human Jumonji gene (JARID2) encoding a protein that regulates transcription and reexpression of a protective fetal program. The Journal of thoracic and cardiovascular surgery 2008, 136 (3): 709–716.

40. Zhong B, Yang Y, Li S, Wang YY, Li Y, Diao F, Lei C, He X, Zhang L, Tien P, Shu HB. The adaptor protein MITA links virus-sensing receptors to IRF3 transcription factor activation. Immunity 2008, 29 (4): 538–550.

41. Li N, Zhou H, Wu H, Wu Q, Duan M, Deng W, Tang Q. STING-IRF3 contributes to lipopolysaccharide-induced cardiac dysfunction, inflammation, apoptosis and pyroptosis by activating NLRP3. Redox biology 2019, 24: 101215.

