## Supplementary material for "Integrated Analysis Reveals Immunogenic Cell Death in Sepsis-induced Cardiomyopathy": figure legend

Figure 1. The immune cell infiltration characteristics during SIC. 
A. Immune cell infiltration characteristics in the mouse hearts at steady state and on days 3/7/21 after CLP. 
B. Immune cell communication characteristics in the mouse hearts at steady state and on days 3/7/21 after CLP.

Figure 2. Monocytes infiltrating to the heart during SIC had shown high ICD activity

Figure 3. Identification of dysregulated ICD-related genes in SIC patients.
A. The heatmap illustrated the expression of significantly upregulated or downregulated ICD-related genes in SIC.
B. Correlation analysis of ICD-related genes, with red and green colors indicating positive and negative correlations, respectively. Correlation coefficients are denoted in the pie chart. There were strong regulatory connections between IFNGR1 and IL6, HSP90AA1 and PRF1, LY96 and IL1R1, as well as IL1R1 and MYD88.
C. Circos plot illustrating ICD-related gene interactions in SIC.

Figure 4. Identifying ICD-Associated Clusters in SIC
A. Matrix displaying consensus clustering results for k = 2. 
B. Cluster-consensus score when k set at different values.
C. Expression patterns of ICD-related genes between the two identified clusters.
D. PCA visualization depicting the distribution of the two identified clusters.

Figure 5. Immune infiltration characteristics between ICD clusters.
A. Comparing the proportions of infiltrated immune cells between the two ICD clusters. 
B. Immune cells infiltration boxplot.

Figure 6. GSVA analysis between ICD clusters.
A GSVA analysis using the GO database. The terms were ranked by t-value.
B GSVA analysis using the KEGG database. The terms were ranked by t-value.

Figure 7. Identification of important modules and hub genes in SIC patients.
A. The scale-free index and mean connectivity at various soft-threshold powers.
B. Dendrogram of differentially expressed genes based on dissimilarity (1-TOM).  The color band indicated module assignments determined by Dynamic Tree Cut.
C. Heatmap of the correlation between module eigengenes and clinical status. The turquoise module was selected as the SIC related module.
D. Scatterplot of gene significance versus module membership in the turquoise module. The strong correlation between GS and MM suggested that the turquoise module was highly correlated with SIC. 

Figure 8. Identification of important modules and hub genes associated with ICD clusters.
A. The scale-free index and mean connectivity at various soft-threshold powers.
B. Dendrogram of differentially expressed genes based on dissimilarity (1-TOM).  The color band indicated module assignments determined by Dynamic Tree Cut.
C. Heatmap of the correlation between module eigengenes and ICD clusters. The purple module was selected as the ICD related module.
D. Scatterplot of gene significance versus module membership in the purple module. The strong correlation between GS and MM suggested that the purple module was highly correlated with ICD. 

Figure 9. Identifying the gene sets for the diagnosis of SIC using machine learning models
A The intersections between SIC-related genes and ICD-related genes from WGCNA analysis.
B Boxplots illustrating the residuals of individual machine learning models, with the red dot indicating the root mean square error (RMSE).
C Cumulative distribution of residuals for each machine learning model.
D The important variables in the SVM model
E Correlation between LVEDD and TCEAL9
F Correlation between LVESD and RARRES1
